# Geometric constraints and cognitive inputs jointly shape emergent brain dynamics and topology

**DOI:** 10.64898/2026.08.07.742341

**Authors:** Ahmad Beyh, Jason Z. Kim, Waheed U. Bajwa, Linden Parkes

## Abstract

How the brain’s physical geometry gives rise to its flexible functional repertoire remains a central question in neuroscience. Here, we approached this question through recurrent neural networks (RNNs). We trained three RNN classes on a cognitive working-memory task, forming a graded hierarchy of spatial constraints: *Vanilla RNNs* (no spatial constraints), *Masked RNNs* (projection constraints limiting where information enters and leaves the network), and *biophysical RNNs* (*bioRNNs*; projection constraints and spatial embedding of the networks’ connectivity using the brain’s inter-regional Euclidean geometry). We assessed how well each RNN class predicted empirical fMRI activity without exposing them to it during training. Our results showed that *bioRNNs* were the only networks to successfully predict empirical brain activity and to organize their dynamics into a spatial pattern that recapitulated the brain’s principal hierarchy (the sensorimotor–association axis). Additionally, *bioRNNs’* ability to predict empirical brain activity emerged along a training trajectory in which the effects of brain geometry were encoded by the network first, then partly traded back as the task was mastered. Importantly, brain-like topological features emerged in *bioRNNs* as they increased their task proficiency while maintaining their ability to predict brain activity. Taken together, our results indicate that physical geometry and cognitive inputs play distinct, complementary roles: while geometry constrains the space of possible brain dynamics, cognitive inputs determine which dynamics are expressed. Our results also situate topology as the scaffold through which the physically embedded brain reconciles wiring costs and computational demands.

## INTRODUCTION

The brain is a physically embedded system^1^. A central goal of neuroscience is to understand how this physical geometry constrains the brain’s complex web of inter-regional connectivity and emergent neural dynamics^2^, as well as how these entwined relationships enable the computational processes that underlie cognition and behavior^3^. Decades of neuroscience research have shown that brain geometry, connectivity, and function are tightly, though imperfectly, coupled^4–10^. While geometry plays a key role in determining the layout of anatomical connections, which in turn shapes the repertoire of dynamics the brain can express^2^, brain function is not a trivial product of either its physical embedding or its connectivity alone^9,11^. Instead, to flexibly respond to complex stimuli, the brain has evolved to selectively uncouple its function from its underlying geometry and connectivity, enabling humans to successfully navigate their ever-changing environment^12^. How this flexible brain function emerges from fixed geometry and connectivity is the focus of this work.

Previous work has shown that the physical embedding of brain regions predicts both their anatomical and functional connectivity, with regions located closer together in space being both more likely to share an anatomical pathway and to exhibit similar dynamics^8,13^. This organization is thought to reflect the brain’s need to produce a cost-efficient solution for its wiring; short-range connections are both metabolically and materially lower in cost than long-range ones^14^. However, long-range connections are well documented in the brain and are thought to enable the efficient communication between distributed systems, which is a prerequisite for behavior^15,16^. Similarly, certain regions, known as hubs, exhibit patterns of strong and diverse structural connectivity that exceed what would be predicted by spatial proximity to their neighbors^17^. These topological properties, among others, are thought to play an important role in explaining how and where brain function diverges from its structure. Thus, while geometry accounts for a substantial fraction of variance in both structural and functional connectivity^6,18^, it is only part of the story^5^.

A flexible mapping between the brain’s geometry, connectivity, and its function is therefore critical to enabling complex human cognition, as it allows the same anatomical substrate to support different activity patterns depending on context, behavioral state, and the presence of sensory information^5,19^. Prior modeling work has sought to characterize this relationship. For example, generative network models that probabilistically add connections over time, absent any task-related learning outcome, have shown that a combination of inter-regional physical distance and topological wiring rules can produce brain-like connectivity, illustrating a relationship between geometry and connectivity^20–22^. Eigenmode decomposition of the cortical sheet has been shown to capture the topography of brain activity maps, illustrating a relationship between geometry and brain function^1^. Or, for example, connectome-based reservoir models, in which empirical structural connectivity is implemented directly as the recurrent weight matrix, can generate neural dynamics and support computation, further linking connectivity to function^23^. In a complementary computational account, large-scale circuit models with hierarchically organized intrinsic timescales have shown how the cortex’s anatomical connectivity can give rise to a functional hierarchy of dynamics, directly linking structure to function within a mechanistic model^24^. However, many such applications (for review, see Suárez et al.^9^) have only captured select components of this relationship piecemeal, *e.g.*, by addressing structure or function separately, and in the absence of behavioral inputs accompanied by corresponding learning goals. Here, we seek to examine the relationship between brain regions’ spatial embedding (brain geometry), connectivity, and function within a single end-to-end model during task learning.

To achieve this goal, we leverage recurrent neural networks (RNNs). RNNs are artificial dynamical systems that can be trained via backpropagation through time to perform a wide range of behavioral and cognitive tasks, and they readily support the integration of geometric constraints into their design^25^. Because their connectivity, task inputs, and dynamics are all fully observable and editable, RNNs can uncover precise details of how and when computational dynamics emerge from connectivity, making them a powerful tool for the causal study of neural computation *in silico*^26–30^. Crucially, recent work has shown that imposing biological constraints, such as spatial embedding and a cost on long-range connections, drives RNNs to converge on structural and functional motifs observed in the cortex^27^. Here, we leverage these insights to develop novel biophysical RNNs (*bioRNNs*) that incorporate biological constraints to study the relationship between brain geometry, connectivity, and function.

In this study, we used *bioRNNs* to investigate how the human brain’s physical geometry constrains the emergence of brain-like neural dynamics and connectivity in these synthetic networks. We do so by constraining the spatial embedding of our RNNs’ hidden-layer nodes while allowing their dynamics and connectivity topology to emerge through task training. By systematically layering spatial constraints onto RNNs and comparing their internal dynamics against empirical human fMRI data, we ask three linked questions: (i) whether spatial embedding allows networks to better reproduce real brain activity; (ii) how spatial embedding and cognitive inputs contribute differentially to emergent brain activity; and (iii) whether brain-like topology supports the acquisition of function and observed dynamics. Our results suggest that physical geometry and structured cognitive inputs are both required to support the emergence of a brain-like topology in *bioRNNs* that captures empirically observed brain activity unseen by the models, and that this capacity to predict activity arises only when *bioRNNs* depart from the fixed geometric scaffold that constrains them.

## RESULTS

### Spatial embedding offers a learning advantage

To assess the impact of geometry on brain structure and function, we developed and trained three classes of RNNs on a delayed match-to-sample with distractors (DMS-D) task^31,32^: *Vanilla RNNs* (**Fig. 1a**), *Masked RNNs* (**Fig. 1b**), and spatially embedded *bioRNNs* (**Fig. 1c**). For each of these RNN classes, we trained 100 randomly initialized RNNs (runs) that each consisted of 100 nodes representing the brain regions of the left hemisphere^33^. *Vanilla RNNs* are standard RNNs whose inputs and internal organization are not shaped by any spatial prior that encodes brain geometry. These represent our baseline RNN. *Masked RNNs* include *projection constraints*, a fundamental neurobiological feature that governs how the brain receives environmental inputs and produces behavioral outputs^34,35^. Here, inputs enter the network exclusively through a subset of nodes representing the visual cortex (**Fig. 1b,c**; *read-in nodes*), while outputs are read exclusively from a different subset representing the association cortex (**Fig. 1b,c**; *read-out nodes*). Finally, *bioRNNs* additionally employ a *spatial embedding* that shapes the network’s inter-node connectivity (*i.e.*, the hidden weights). Throughout training, these weights are shaped by inter-regional Euclidean distances derived from empirical brain data, thereby prioritizing a connectivity pattern that mirrors the brain’s geometry. Thus, our RNNs form a graded hierarchy of increasing geometric constraints, spanning from no constraints (*Vanilla RNNs*) to simple brain-like constraints on inputs and outputs (*Masked RNNs*), to ones that also encode how the brain’s geometry influences its connectivity (*bioRNNs*). See *RNN overview* section in Materials and Methods for more details.

**Figure 1.**
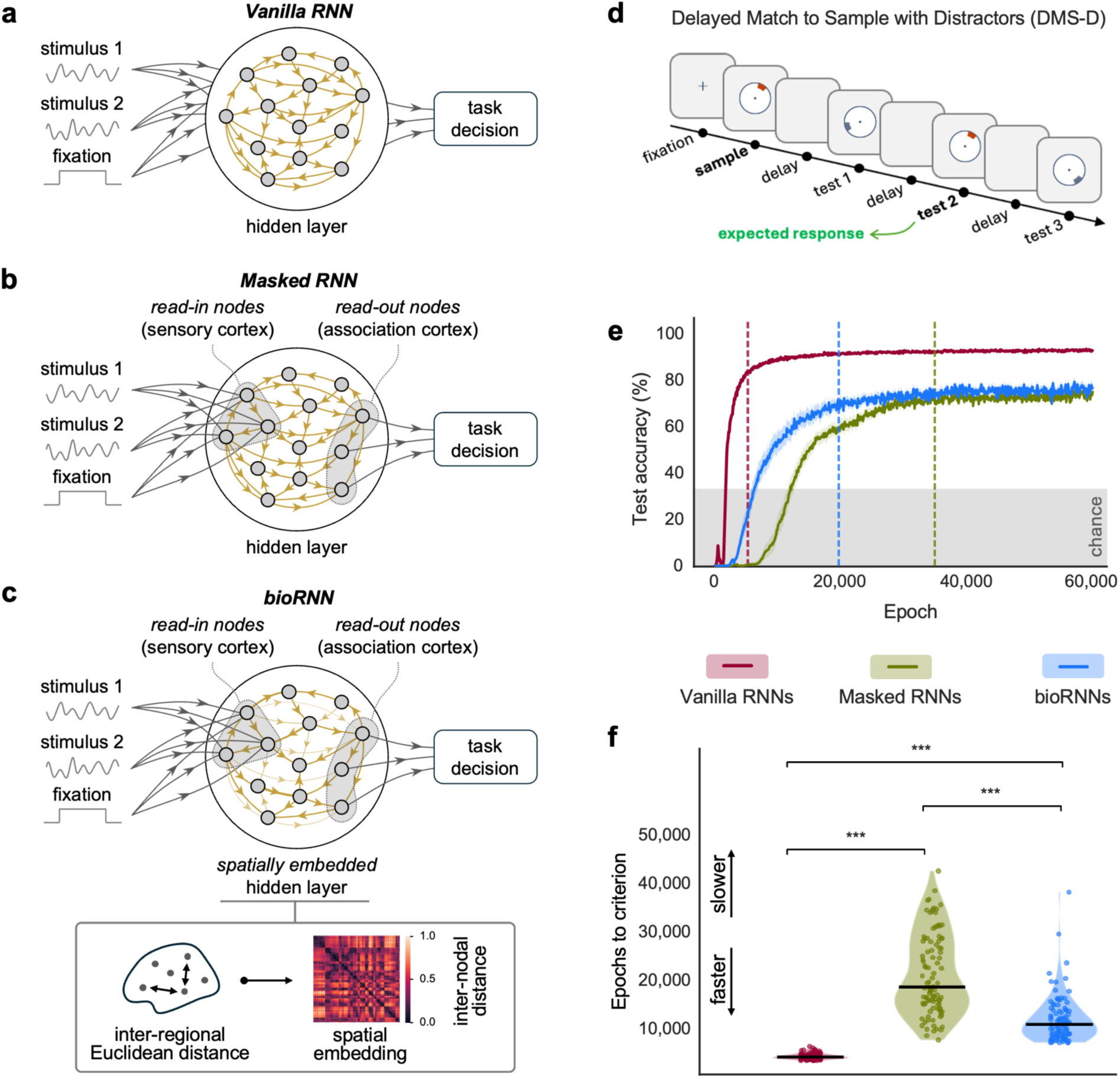
Spatial embedding offers *bioRNNs* a learning advantage in a working-memory task. We trained three RNN classes on a delayed match-to-sample with distractors (DMS-D) task akin to an N-back working-memory task. **(a)** *Vanilla RNN*: a fully recurrent network that receives the task inputs at all hidden units and reads the task decision from all units, with no spatial priors. **(b)** *Masked RNN*: identical to the vanilla network except that inputs enter only through a fixed subset of *read-in nodes* (here, visual cortex) and outputs are read only from a separate subset of *read-out nodes* (here, association cortex), imposing *projection constraints* as the only spatial prior. **(c)** In *bioRNNs*, a spatial embedding is added to the masked architecture, in which the hidden-to-hidden weights are shaped throughout training by a neurobiological prior. Here, we use the brain’s inter-regional Euclidean distance matrix to constrain the relative spatial patterning of connectivity during RNN training. Thus, *bioRNNs* impose two interrelated spatial priors, one based on task inputs and outputs and another based on brain geometry. **(d)** We trained RNNs on a DMS-D task^31,32,38^. A trial began with a fixation cue and a sample stimulus, after which three test stimuli were presented in sequence, separated by delays; the network was to respond only to the test that exactly matched the sample. **(e)** Test accuracy across training (mean ± 95% confidence interval over 100 networks, or runs, per RNN class). All classes successfully learned to perform the task well above the 33% chance level, but they did so at different rates. The dashed vertical lines captured when each RNN reached the training criterion, *i.e.*, when 95% of the runs reached their accuracy asymptote (see **Fig. S1**). **(f)** Violin plots of the number of training epochs to the criterion for each class. *bioRNNs* reached the performance criterion faster than *Masked RNNs* despite having more stringent training demands. Statistical significance is based on Wilcoxon signed-rank test.

The DMS-D is a complex working-memory task that engages temporal integration, perceptual decision-making, and inhibitory control, all key facets of higher-order cognition^36,37^ in humans, making it well-suited to probing the neural dynamics that underlie complex behavior. We trained our RNNs on task data simulated using NeuroGym^38^. Each DMS-D task trial began with a brief fixation signal indicating the start of a new trial, followed by the presentation of a unique target stimulus consisting of 32 simultaneous inputs (**Fig. 1d**). Three additional test stimuli were then presented in sequence, separated by delays, and the RNN was required to respond only when it encountered a test stimulus that was an exact match to the initial target stimulus. As such, the RNN’s decision relied on maintaining a representation of the target stimulus in working memory throughout the trial and inhibiting its response until an exact match was encountered.

We found that all RNN classes successfully learned the task (**Fig. 1e**). Specifically, given that the DMS-D task required correctly identifying one of the three test stimuli as the correct match, chance performance was set at 33% accuracy. All models exceeded this chance baseline. While *Vanilla RNNs* reached the highest final accuracy (92.00 ± 1.52%), *Masked RNNs* and *bioRNNs* achieved similar but lower levels (74.26 ± 6.25% and 75.92 ± 10.23%, respectively). This result suggests that minimally constrained *Vanilla RNNs*, which were only optimized to solve the task, remained the best performers. There were also clear differences in learning speed between the RNNs. We fit each RNN’s learning curve with a logistic function (**Fig. S1**) and assessed when task performance stabilized (*i.e*., reached 95% of its asymptote). Although we trained the RNNs up to 60,000 epochs, we observed that all RNN classes reached stable task performance within 40,000 epochs. Therefore, we used this marker as the anchor for all subsequent analyses. Here, we observed that *Vanilla RNNs* were the fastest to learn the task (4,019 ± 615 epochs; matched-pairs rank-biserial correlation *r_rb_* = -1.0, Wilcoxon signed-rank *p* < 0.001 against both other RNN classes), consistent again with the idea that a minimally constrained RNN optimized for performance rather than biological relevance is best positioned to learn most rapidly (**Fig. 1f**). Critically, however, *bioRNNs* learned significantly faster (11,729 ± 4,834 epochs) than *Masked RNNs* (21,150 ± 8,348 epochs; *r_rb_* = -0.85, *p* < 0.001). This was the case despite the fact that the spatial penalty imposed on *bioRNNs* during training posed the largest burden among the three RNN classes (see spatial loss and task loss evolution in **Fig. S2**). Therefore, in the presence of *projection constraints*, a fundamental property of how visual inputs project to the brain, *spatial embedding* afforded the networks a significant learning-rate advantage. This observation is consistent with prior reports showing that biologically grounded physical constraints can serve as inductive biases that enhance task learning, rather than just as additional costs^27^.

### *bioRNN* dynamics capture empirical brain dynamics and the cortical hierarchy

Having established that the brain’s spatial embedding confers a learning advantage in the presence of biologically-informed *projection constraints*, with a minimal impact on task performance, we next assessed how well our RNNs captured real, empirical brain activity observed *in vivo* in humans using functional MRI (fMRI) data obtained from 100 healthy young adults (Human Connectome Project; HCP-YA)^39^. First, in order to isolate the dominant, behaviorally relevant modes of high-dimensional activity^40^, we applied principal component analysis (PCA) to the time series of the fully trained RNNs to capture their hidden-state trajectories in a low-dimensional subspace (**Fig. 2a**). The first five resulting PCs captured 72.7% ± 1.2% of hidden-activity variance in *Vanilla RNNs*, 77.2% ± 2.2% of hidden-activity variance in *Masked RNNs*, and 76.2% ± 3.5% of hidden-activity variance in *bioRNNs* (see **Fig. S3** for more details on PC selection). Importantly, projecting the weights of the PCs onto the brain surface did not yield spatially smooth PCs for either *Vanilla RNNs* (Moran’s *I* = -0.01 ± 0.01) or *Masked RNNs* (Moran’s *I* = -0.01 ± 0.01), but a spatially smooth organization did emerge for *bioRNNs* (Moran’s *I* = 0.18 ± 0.06), indicating that the spatial embedding of the hidden layer pushed the *bioRNNs* to organize their activity in a brain-like pattern (**Fig. 2b**). In other words, once the recurrent layer is encouraged to reflect cortical geometry, its dominant activity modes recapitulate smooth spatial gradients like those observed in large-scale cortical fMRI patterns^41,42^.

**Figure 2.**
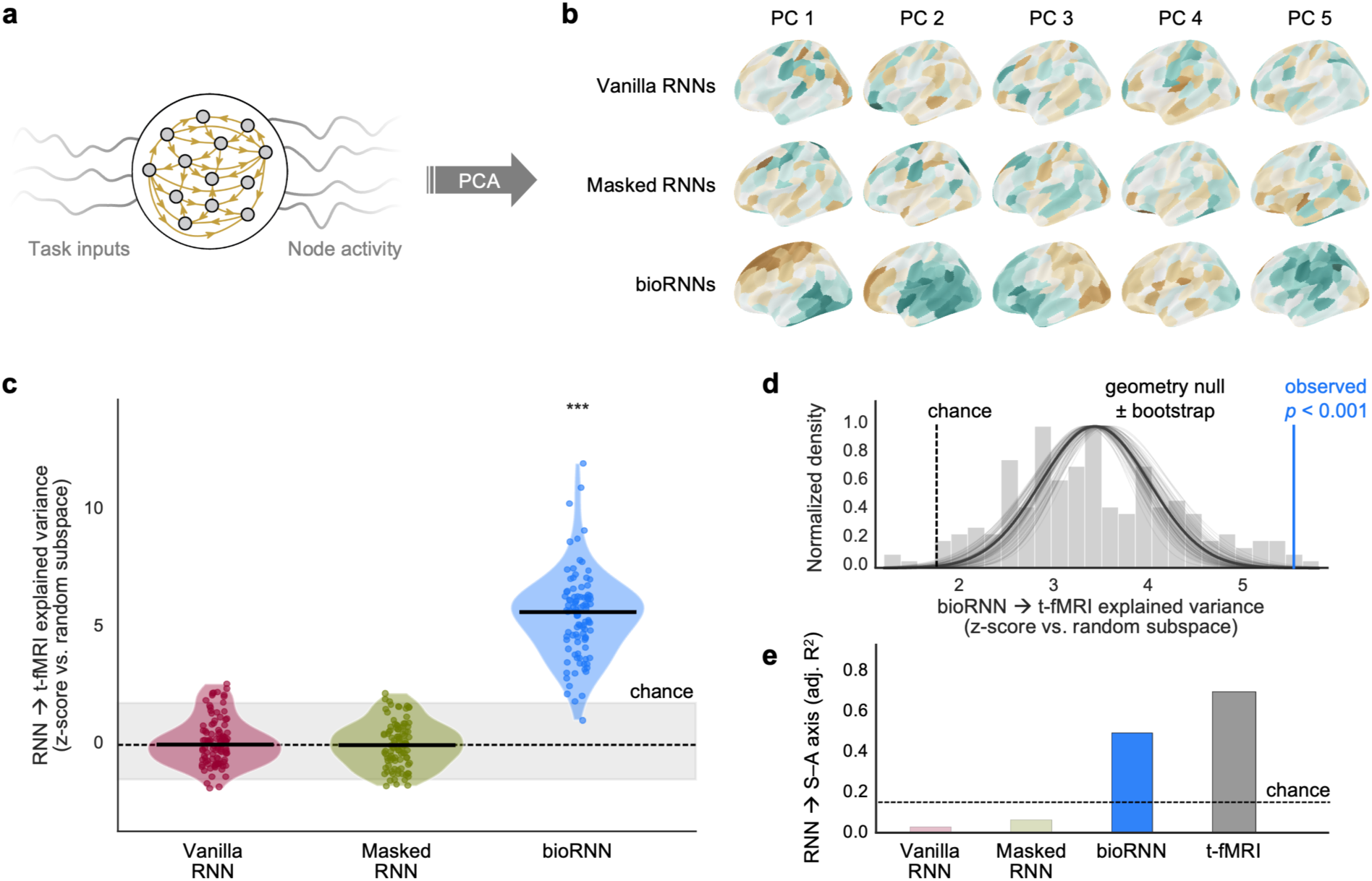
bioRNN dynamics capture empirical fMRI dynamics and the cortical hierarchy. **(a)** Each trained RNN’s hidden-state time series was transformed to a low-dimensional representation through principal component analysis (PCA). **(b)** Weights of the first five principal components (PCs) projected onto the cortical surface. *Vanilla RNNs* and *Masked RNNs* showed no discernible spatial structure, whereas *bioRNNs* displayed smooth, spatially organized PC maps. **(c)** Variance in the N-back task fMRI dynamics explained by the first five PCs of each RNN class, expressed as a z-score relative to a random-subspace baseline (*z_sub_*; grey band, chance). Only *bioRNNs* exceeded chance. **(d)** *bioRNNs* were retrained with the spatial embedding permuted to preserve its spatial autocorrelation while breaking its correspondence to the cortex (grey histogram, with bootstrap density curves obtained through linear mixed-effects models). The veridical embedding (red) exceeded the entire null distribution (*z_geom_* = 3.67, *p* < 0.001; 95% CI [2.76, 5.21]), indicating that alignment with empirical dynamics depends on the true anatomical arrangement of nodes rather than on spatial smoothness alone. **(e)** Variance in the cortical sensorimotor–association (S–A) axis explained by each RNN class and by empirical task fMRI PCs (dashed line: chance). Only *bioRNNs* recovered the S-A axis (adj. R^2^ = 0.49) and do so to a moderately close degree as task fMRI itself (adj. R^2^ = 0.68).

Next, we used the first five RNN PCs as predictors of empirical fMRI activity derived from the N-back working-memory task^43^. Because even a randomly chosen five-dimensional subspace captures a non-trivial fraction of variance in high-dimensional fMRI data, raw explained variance cannot be interpreted on its own^44,45^. We therefore benchmarked each RNN against a null of 1,000 random five-dimensional subspaces and expressed its fMRI explained variance as a z-score relative to this random-subspace baseline (*z_sub_*; see *fMRI variance explained and the random-subspace* in the Methods). We observed that *bioRNNs* were the only ones that exceeded chance in predicting variance in fMRI data (*z_sub_* = 5.54 ± 1.87, *p* < 0.001; **Fig. 2c**). Meanwhile, *Vanilla RNNs* and *Masked RNNs* yielded *z_sub_* values around zero, indicating that their ability to predict empirical brain activity was no better than chance. This result shows that neither unconstrained RNNs (*Vanilla*) nor those with segregated inputs and outputs (*Masked*) were sufficient to generate cortex-like neural dynamics. Rather, what matters is the specific spatial arrangement of the recurrent connectivity engendered by brain geometry. Notably, our *bioRNNs* were trained on a synthetic DMS-D task *in silico* yet successfully predicted N-back fMRI time series acquired independently in humans *in vivo*. This is a striking form of transfer, in which *in silico* networks trained only on a synthetic task predicted the low-dimensional representation of the moment-to-moment dynamics of living human brains performing a conceptually similar but distinct working-memory task without ever encountering that data during training. Finally, we observed this correspondence at the individual-subject level as well: *bioRNNs* whose hidden weights more closely resembled a subject’s inter-regional Euclidean geometry better predicted that subject’s fMRI activity (*r_S_* = 0.25 ± 0.09, *p* < 0.001; **Fig. S4**).

An important question emerged from our previous results: can *bioRNNs’* prediction of empirical fMRI dynamics be attributed solely to the spatial autocorrelation introduced by the Euclidean embedding, rather than to the veridical arrangement of nodes in anatomical space? To address this, we constructed a geometry null in which the embedding was permuted to preserve its spatial autocorrelation while breaking its correspondence to cortex^13,46^, holding the task and *projection constraints* fixed, and fully retrained 200 such RNNs up to 40,000 epochs (the training criterion established previously). Spatial autocorrelation accounted for part of the effect: the 200 null RNNs predicted fMRI dynamics above the random subspace baseline (mean *z_sub_* = 3.42 ± 0.91; **Fig. 2d**). Critically, however, the predictive performance of the veridical *bioRNN* significantly exceeded the null distribution (*z_geom_* = 3.67, *p* < 0.001; 95% CI: *z_geom_* = [2.76, 5.21], *p* = [<0.001, 0.003]; **Fig. 2d**). Thus, the *bioRNNs’* alignment with empirical brain activity depends on the veridical spatial embedding and its congruence with the cortical locations the nodes represent, beyond what spatial autocorrelation alone can produce. In other words, in addition to spatial smoothness, the particular pattern in which the cortex is laid out in space endows these networks with brain-like dynamics, demonstrating that the anatomical positions of regions relative to one another, and how these positions interact with the *projection constraints*, carry functionally relevant information.

Given the ability of *bioRNN* dynamics to predict unobserved functional brain activity, we next asked how well these dynamics aligned with the brain’s known hierarchical organization, which we summarized using the sensorimotor–association (S–A) axis^47^. Specifically, we ran regression analyses in which the RNN PC weights acted as predictors and the S–A axis as a target variable. We found that *bioRNNs* were the only ones whose simulated dynamics captured the cortical hierarchy (adj. R^2^ = 0.49; **Fig. 2e**). Here as well, this ability was not the simple product of the spatial autocorrelation present in both the *bioRNNs* and the S–A axis, which we confirmed with the same null-testing approach we used in the dynamics prediction analysis (*p* = 0.002). Meanwhile, *Vanilla RNNs* and *Masked RNNs* captured nearly none of the hierarchy (adj. R^2^ = 0.04 for both). For comparison, we ran an additional regression analysis that used PC weights derived directly from N-back task fMRI activity as predictors and found that task fMRI activity mapped onto the S–A axis moderately better than *bioRNNs* did (adj. R^2^ = 0.68; **Fig. 2e**), and this was also not solely attributable to spatial autocorrelation (*p* = 0.001). Thus, *bioRNN* dynamics explain the dominant hierarchical brain axis to a level approaching the ceiling imposed by fMRI, indicating that they successfully capture a large-scale organizing principle of the cortex. Strikingly, *bioRNNs* recover this cardinal axis of cortical organization without ever being directly exposed to it, whereas fMRI is one of the features used to construct the S–A axis. This result implies that a spatial geometry prior, once shaped by task demands into connectivity, is able to reconstruct the brain’s principal functional hierarchy.

### Working memory training reorganizes *bioRNN* dynamics toward human cortical activity

Having established that *bioRNNs* spatially embedded with the brain’s Euclidean distance structure were the only RNNs that predicted empirical brain activity, we next asked how the similarity between *bioRNNs’* hidden weights matrix and the brain’s Euclidean distance matrix evolved over training. Motivated by the possibility that a network must first acquire brain-like structure before it can express brain-like function^48^, we further asked how the evolution of this similarity relates to *bioRNNs*’ task accuracy and their ability to predict empirical brain activity. To this end, we plotted fMRI explained variance (*z_sub_*) as a function of the alignment (cosine similarity) of the *bioRNN* hidden weights matrix with the brain’s empirical Euclidean geometry, color-coding the resulting trajectory by task accuracy (**Fig. 3a**). Rather than a monotonic increase in both quantities, the trajectory traced a pronounced hysteresis loop with three features that were consistent across networks, and that unfolded in a stage-like manner. In the first stage (**Fig. 3a**, i), structure preceded function: *bioRNN*–brain structural similarity rose steeply early in training, while fMRI prediction remained near chance, and accuracy remained at 0%. In the second stage (**Fig. 3a**, ii), fMRI alignment rose sharply and largely preceded the gain in behavioral accuracy. In the third and final stage (**Fig. 3a**, iii), *bioRNN*–brain similarity reversed course and reduced, such that the training end-state occupied an intermediate spatial similarity regime that maintained a relatively high fMRI prediction alongside high task accuracy. Consequently, a given *bioRNN*–brain structural similarity value maps onto markedly different levels of fMRI prediction, depending on the training phase. For example, the gray band in **Fig. 3a** highlights two points along the training trajectory where *bioRNNs* shared the same moderate level of structural alignment to the embedding (*cosine* ≈ 0.25), but showed markedly different fMRI predictions. That is, *bioRNNs* early in training did not exceed chance in predicting fMRI activity (*z_sub_* ≈ 1.5), but the same *bioRNNs* later in training achieved predictions that far exceeded chance (*z_sub_* ≈ 6.3) despite sharing the same structural alignment with the brain’s Euclidean geometry. Brain-like dynamics in *bioRNNs* were therefore not a fixed function of how Euclidean-like the network geometry was, but were acquired along a training trajectory in which geometry was laid down first and partly traded back as the task was mastered. Together, these findings support a view of the brain in which brain geometry is necessary but not sufficient for task learning, and in which eventual mastery of cognitive processing requires a flexible departure of brain function from the underlying geometry. We also replicated this effect using a Spearman similarity metric to confirm that it is not simply a product of metric choice (**Fig. S5**).

**Figure 3.**
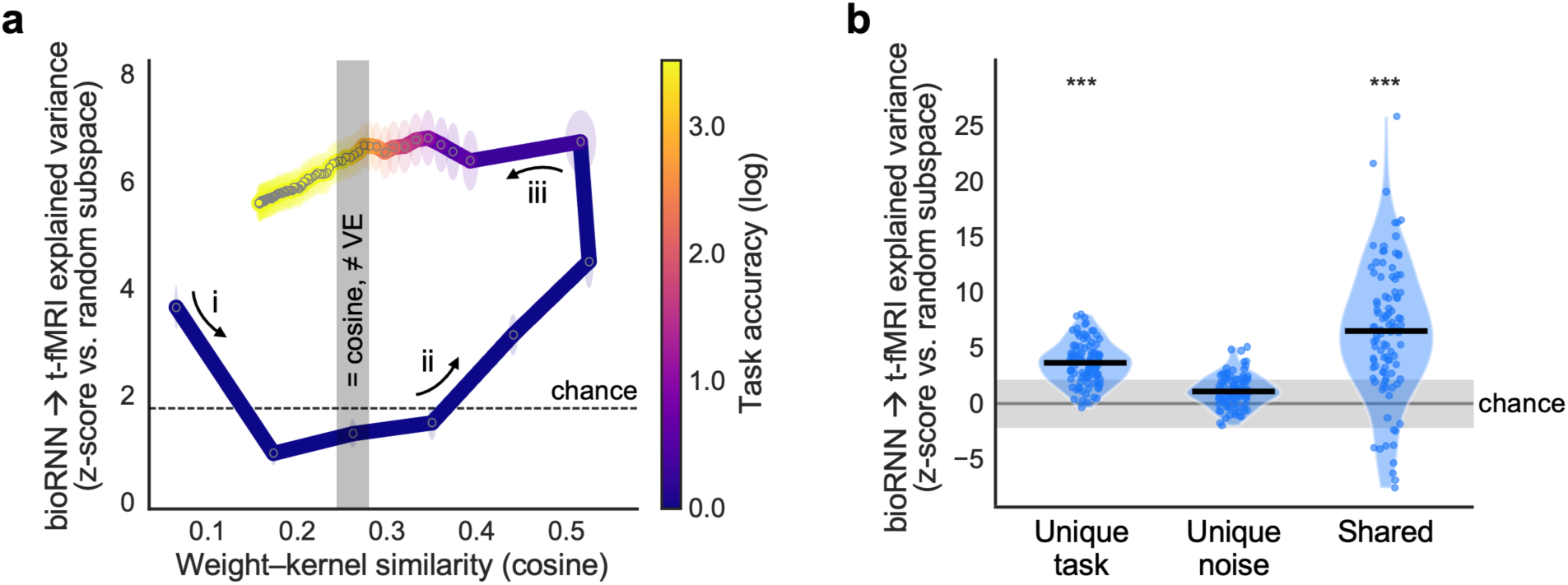
Task learning shapes emergent dynamics. **(a)** Training trajectory of *bioRNNs* as a function of fMRI explained variance (y-axis; *z_sub_*) versus weight–kernel similarity (x-axis; cosine similarity between the hidden-weight matrix and the brain’s Euclidean distance kernel), with points colored by task accuracy (log scale). The trajectory formed a hysteresis loop with three stages: (i) structural similarity rose (rightward shift along the x-axis) while functional alignment remained near chance; (ii) functional alignment rose steeply (upward shift along the y-axis), still at high structural similarity, largely preceding the gain in accuracy (dark purple coloring); and (iii) structural similarity dropped to an intermediate value while functional alignment remained high and task accuracy increased (yellow coloring). The oval-shaped disks at each point along the trajectory represent the 95% CI along both axes. The gray band highlights two positions along the trajectory that share the same weight–kernel similarity (cosine ≈ 0.25) but with markedly different fMRI predictions (*z_sub_* ≈ 1.5 vs. *z_sub_* ≈ 6.3). **(b)** Variance decomposition analysis based on fully-trained *bioRNNs* tested with structured task inputs and with random noise inputs. The unique-task and shared (task and noise together) components predicted fMRI activity above chance, whereas the unique-noise component did not. Interestingly, the shared basis was the strongest predictor.

To further isolate the unique contributions of geometry and structured task inputs in shaping emergent dynamics, we tested *bioRNNs* with unstructured noise inputs. By feeding noise into the trained networks, we assessed the emergent dynamics that their structure supports, regardless of the temporal structure of their inputs. We then performed a variance decomposition to assess the unique and shared contributions of each regime (*bioRNN_task_* and *bioRNN_noise_*) to the prediction of empirical fMRI activity (see *Variance decomposition of task-evoked and intrinsic dynamics* in Materials and Methods). Using singular-value decomposition, we constructed a basis of pooled predictors from the sets of PCs derived from *bioRNN_task_* and *bioRNN_noise_*. We then asked how much unique fMRI variance each regime explained compared to the shared basis that captured the variance common to both the task-driven and noise-driven regimes. Here, the shared component captures geometry’s effect, whereas the unique task and noise components isolate the effect of the task’s temporal structure.

The results (**Fig. 3b**) revealed that the task-driven *bioRNN* dynamics contained unique information predictive of fMRI activity that the shared basis did not contain (mean *z_sub_* = 3.52 ± 1.84, *p* < 0.001 based on a Wilcoxon signed-rank test against the subspace chance upper bound), but noise-driven dynamics were not uniquely predictive (*z_sub_* = 1.07 ± 1.37, *p* = 1.00). Interestingly, the shared basis was the strongest predictor of fMRI activity (*z_sub_* = 6.25 ± 5.93, *p* < 0.001). This result suggests that most of the *bioRNNs’* prediction of empirical brain activity is carried by the shared component, *i.e.*, the dynamics common to the task-and noise-driven regimes. Because this signal emerges whether the network is driven by structured input or by unstructured noise, it is most plausibly generated by the intrinsic dynamics of the network’s spatial embedding (geometry) rather than by the input itself. However, on top of this shared, architecture-driven baseline, task input contributed additional, uniquely predictive dynamics, whereas unstructured noise did not. Together, these results indicate that the embedded geometry establishes a baseline correspondence with cortical activity, while structured task input is required to capture the aspects of empirical dynamics that geometry alone cannot produce.

### *In silico* evidence for topology’s functional relevance

The third stage of the hysteresis trajectory (**Fig. 3a**) suggested that continued refinement of hidden-layer connectivity within *bioRNNs* supported task learning and enabled them to maintain explanatory power of empirical brain dynamics despite departing from a simple Euclidean embedding. In this final section, we sought to explain this departure by examining network topology.

The topology of the human connectome is not random: it interleaves locally clustered, segregated communities with a small set of highly connected hubs that integrate information across the brain, an organizational signature of mammalian brains thought to underpin efficient neural communication and flexible cognition^11,17^. Critically, this complex topology is not predicted solely from brain geometry, as a network designed to minimize wiring cost alone would tend toward a regular, lattice-like organization, whereas the empirical connectome interleaves short-range clustering with a minority of costly long-range links that support global integration^7,49^. We therefore reasoned that the departure of our *bioRNNs* from geometry with task learning may reflect the emergence of brain-like topology in their underlying connectivity.

To investigate this possibility, we assessed the topology of the intermediate hidden-layer nodes (*i.e.*, nodes not labeled as *read-in* or *read-out*) during the final learning stage, which is marked by rapid task learning. We chose these nodes to examine topology because they fall outside the direct influence of the *projection constraints* on the overall network structure, which necessarily induces a tripartite structure within the networks (**Fig. S6**). Assessing the functional connectivity within the *bioRNN* intermediate subnetwork, we observed patterns suggesting the presence of modules resembling those observed in empirical functional connectivity (**Fig. S7**). Given the known role topology plays in supporting learning and dynamics, we assessed its emergence in both *bioRNNs* and *Masked RNNs*. Note, because *Masked RNNs* lagged *bioRNNs* in task learning, we aligned the topology trajectories of the two RNN classes to facilitate comparison, and this required a 6,000-epoch shift (**Fig. 4**). Topological changes along the full training trajectory are reported for *bioRNNs* in **Fig. S8** and for *Masked RNNs* in **Fig. S9**.

**Figure 4.**
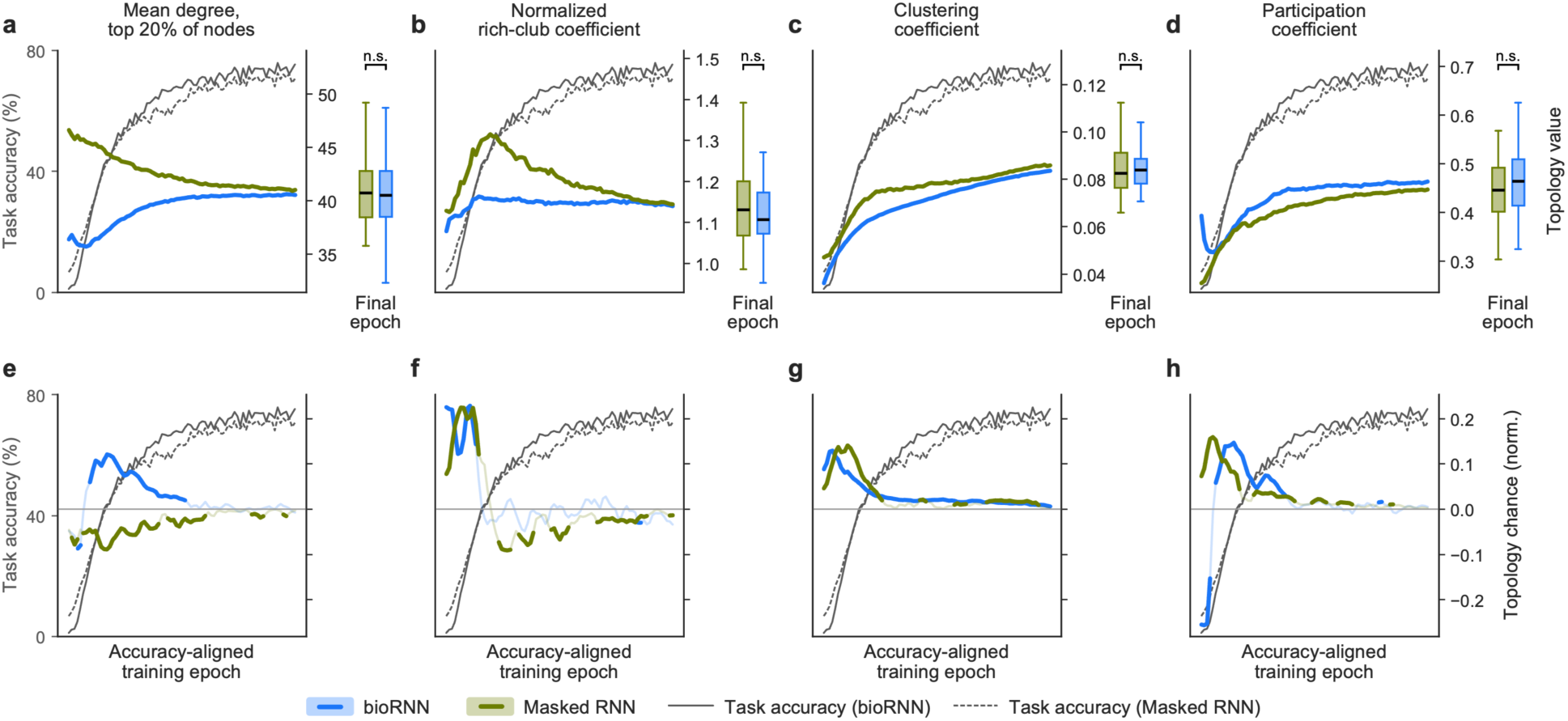
Connectome-like topology emerges during training and tracks task learning. Graph-theoretic properties of the intermediate (non-projection) nodes of *bioRNNs* and *Masked RNNs* across training, with task accuracy (black) overlaid on each panel. The trajectories of the two RNN classes were aligned by shifting the *Masked RNN* traces by 6,000 epochs before plotting to facilitate a direct comparison. **(a)** The mean degree of the top 20% of nodes captured the emergence of strongly connected nodes (hubs). **(b)** The normalized rich-club coefficient converged on a value >1.00, indicating the formation of a densely interconnected hub core. **(c)** The mean clustering coefficient increased, reflecting the emergence of local, segregated processing units. **(d)** The participation coefficient also increased with training, reflecting increasingly diverse cross-module connectivity (integration). **(e)** The mean connection length of the top 20% of edges (by length) rose sharply immediately before the *bioRNNs* learned the task, indicating that the departure from simple brain geometry was driven by changes favoring the selective development of long-range connections. The length analysis does not apply to *Masked RNNs*. **(f–j)** Rate of change of each metric as a function of training, showing that topological reorganization was concentrated in the early part of task learning. See **Fig. S8** and **Fig. S9** for the full trajectory of each RNN class.

Our analysis revealed that two neurobiological topology features emerged in the intermediate hidden-layer node of both *bioRNNs* and *Masked RNNs*, but did so following markedly different trajectories. First, the mean degree of the top 20% of nodes (**Fig. 4a**) and the normalized rich-club coefficient (**Fig. 4b**) showed that both RNN models developed hub nodes that were preferentially interconnected. However, these features developed steadily before reaching a stable plateau in *bioRNNs*, whereas *Masked RNNs* initially overshot the *bioRNNs*’ trajectory early, before converging toward it later, settling on a similar final value at the end of training. Furthermore, this pronounced reduction in topology in *Masked RNNs* coincided with a reduction in the relatively small amount of fMRI activity they could explain early in training (**Fig. S10**).

Second, the clustering coefficient (**Fig. 4c**) and participation coefficient (**Fig. 4d**) revealed that both RNNs simultaneously increased their local clustering and their cross-module communication over training. In other words, the networks became both more segregated, forming locally dense and specialized neighborhoods, and more integrated, distributing their connections across communities to enable cross-talk. Here, there was a subtler difference between the two RNN classes: during training, *bioRNNs* formed slightly fewer clusters that were more integrated (higher participation coefficient) than *Masked RNNs*, but this difference was insignificant by the end of training. The increased integration in *bioRNNs* was supported by the development of long-range connections within the networks (**Fig. S8**). Importantly, these long-range connections emerged despite the high cost imposed by the embedding. In *bioRNNs*, the spatial embedding does not explicitly penalize long-range connections, but constrains the relative spatial patterning of connectivity; the spatial loss is minimized when recurrent weights decrease with interregional distance. Thus, the emergence of long-range connectivity prior to task learning supports their critical functional role.

Finally, the largest changes across all metrics occurred early, right around the initial rise in task accuracy, and gradually stabilized as accuracy plateaued (**Fig. 4f–j**). This result indicates that the network’s topological architecture was largely established early, before the task was mastered, but required fine-tuning to support task learning. Topological reorganization, therefore, preceded behavior rather than following it, mirroring the structure-before-function ordering seen in the hysteresis trajectory. In turn, our results suggest that acquiring a brain-like topology plays an important role in task learning, rather than simply resulting from it^48^. Critically, taken together, our results show that only *bioRNNs* gave rise to sustained brain-like topology that simultaneously enabled task mastery as well as the highest levels of fMRI activity prediction.

## DISCUSSION

How the brain’s physical geometry gives rise to its flexible functional repertoire is a central question in neuroscience. To address this question, we built spatially embedded RNNs that learn a cognitive task under neurobiological constraints and asked how geometry, connectivity, and function interrelate during learning. Our results show that an interaction between the brain’s spatial embedding and its cognitive inputs is critical for the emergence of brain-like task-based dynamics. By layering spatial constraints onto RNNs and allowing their topology and dynamics to emerge during training, we showed that *bioRNNs* could accurately predict empirically observed brain activity out-of-sample, and that this predictive ability was preceded by the acquisition of stable connectome-like topology. Four specific findings emerged. First, brain geometry conferred a learning advantage over circumscribed projections of task inputs and outputs to canonical brain systems. Second, only spatially embedded networks predicted empirical fMRI activity and recovered the principal axis of cortical organization, and this depended on the veridical arrangement of regions rather than solely on the spatial autocorrelation inherent to the underlying RNN architecture. Third, the correspondence with brain activity was acquired along a path-dependent trajectory in which geometry was laid down first and partly traded back as the task was learned. Fourth, without explicit optimization for it, the networks developed a connectome-like topology prior to mastering the task and sustained it thereafter.

Spatial embedding was necessary for any correspondence with empirical brain dynamics. Only *bioRNNs* exceeded the random-subspace baseline in predicting variance in fMRI activity, while *Vanilla* and *Masked RNNs* performed at chance levels. Critically, this advantage was not the simple product of the spatial autocorrelation that our distance-based learning kernel necessarily imposes on the hidden layer. When we tested *bioRNNs* against a geometry null in which the embedding was permuted to preserve spatial autocorrelation while breaking the correspondence with cortical locations, the null RNNs retained part of the effect, but the veridical embedding’s performance exceeded the null distribution. The correspondence, therefore, depended on the specific arrangement of nodes in anatomical space, not on spatial smoothness alone, consistent with the proposal that wiring-cost constraints are a fundamental organizing principle of neural systems^2,7,14^. More broadly, these results support previous work showing that the spatial layout of the cortex plays a critical role in shaping the large-scale activity patterns that the brain can express^1^.

Spatial embedding also organized the networks’ dynamics along the brain’s principal functional hierarchy. The leading principal components of *bioRNN* activity, but not those of *Vanilla* or *Masked RNNs*, recovered the S–A axis, and did so to a degree approaching that of empirical task fMRI. This is notable because the S–A axis was never supplied to the RNNs; it emerged from the interaction between a purely distance-based connectivity prior and task training. Cortical geometry, therefore, appears to confer the capacity for hierarchical organization, which experience then realizes. We additionally observed a subject-specific effect: the more a *bioRNN*’s internal structure resembled a particular brain, the better it predicted that brain’s dynamics. This indicates that the structure–function mapping recovered by these RNNs is not solely a group-average phenomenon. A natural next question is how this experience-dependent hierarchy relates to the molecular, laminar, and transcriptomic gradients to which the S–A axis is highly correlated, gradients that are themselves laid down under tight spatial and metabolic constraints during development^41,47,50,51^. Rather than competing explanations, geometry and molecular patterning may represent coupled expressions of a shared developmental cost structure, a hypothesis that future *bioRNNs* seeded with empirical gradient maps could test directly.

The relationship between *bioRNN* constraints, task learning, and empirical brain activity revealed a multi-stage process of task mastery. The training trajectory traced a pronounced hysteresis loop in which structure was laid down before function: *bioRNN*–brain structural similarity rose steeply early in training, while fMRI prediction remained near chance. Prediction of empirical activity then rose sharply and largely preceded the gain in behavioral accuracy, after which structural similarity decreased to an intermediate level that both preserved high explained variance of fMRI data and allowed for task learning. Brain-like dynamics were therefore not merely a fixed function of how brain-like the network geometry was: the same structural similarity value mapped onto markedly different functional predictions depending on the training stage. Thus, in our model, geometry sets the stage for brain-like dynamics but, by itself, does not produce them, suggesting that geometry defines the space of possible dynamics while experience selects which of those dynamics are realized.

Our variance decomposition analysis made the contribution of task input explicit. Task-evoked *bioRNN* dynamics carried unique predictive information about empirical fMRI activity that the shared basis did not contain, whereas noise-driven dynamics carried no unique predictive information. The shared basis was nonetheless the strongest single predictor, indicating that much of the predictive signal is common to both regimes and is attributable to the embedded architecture itself. Intrinsic, structurally determined dynamics thus supply a substantial baseline correspondence with cortical activity, but the additional component that distinguishes empirical task dynamics appears only when the network is driven by task input; unstructured noise passing through the same architecture does not recover it. This echoes the role of input-driven recurrent dynamics in shaping context-and task-dependent neural activity^29,32^. This result also posits cortical dynamics as a two-tiered phenomenon. First, they are predominantly shaped by geometry, which constrains the space of possible emergent dynamics. Second, interactions with the environment (*e.g.*, via task inputs) are a necessary complement and form a second tier that captures unique aspects of those dynamics.

Beyond simple cortical geometry, the topology that emerged over training tracked each network’s task learning. Throughout training, *bioRNNs* developed a preferentially connected hub core, evident as rising node degree and a rising normalized rich-club coefficient, together with increasing clustering and participation, and a selective development of long-range connections. In other words, *bioRNNs* acquired both local segregation and cross-module integration, which are defining features of small-world cortical organization^17,52,53^. Crucially, these features emerged most rapidly during the earliest phase of learning and then stabilized, indicating that a cortex-like topology was laid down before task mastery rather than arising as its consequence. This ordering mirrors cortical development, in which hub regions and rich-club connections are established early and subsequently refined^54,55^. Our results, therefore, suggest that the brain’s hub-and-rich-club scaffold can be laid down before, and in the service of, the functions it is later refined to support. Importantly, our spatially constrained networks recovered these canonical cortical wiring motifs without explicit optimization for them, further bolstering the idea that geometric wiring constraints strongly shape the connectivity patterns that can emerge in the brain^2,7,27^. Notably, although similar topological features also emerged in *Masked RNNs*, they did not track task learning like they did in *bioRNNs*, and they developed along a less stable trajectory. Moreover, these topological features were accompanied by a selective strengthening of longer connections in *bioRNNs*, allowing them to maintain their ability to predict empirical brain activity while mastering the task. These results suggest that brain geometry and projection constraints interact to drive the specific pattern of cortical organization during learning. Therefore, what makes a system predictive of empirical brain activity is not only its final network-level topological features, but also the trajectory it takes to reach those features, and the interaction between task demands and physical constraints along that trajectory. This also speaks to the many-to-one problem that governs network systems, whereby many configurations can lead to similar topological features, but this does not necessarily lead to dynamics that are more brain-like unless the emergence of these features is shaped by the right constraints during training.

More broadly, our results point to the necessity of importing brain structure into computational modeling tools. They also make a strong case for the potential of *bioRNNs* as *in silico* testbeds for counterfactual experiments, for asking “what if” questions about brain structure and function that cannot be investigated directly in the living brain. For example, one could silence a specific projection, rescale the wiring-cost penalty, or substitute one individual’s geometry for another’s and read out the causal consequences for dynamics and topology. These experiments would extend connectome-based reservoir and constrained-RNN approaches^23,27^ from fixed to manipulable architectures. By independently manipulating spatial embedding, input structure, and connectivity, *bioRNNs* offer future studies a tractable platform for causally probing how anatomy and experience jointly give rise to neural dynamics.

Several limitations are worth considering. First, our RNNs were evaluated against BOLD fMRI data, an indirect and temporally coarse proxy for neural activity; the correspondence we report is between low-dimensional summaries of RNN hidden states and of BOLD dynamics, not between RNN units and spiking activity. Nevertheless, the low-dimensional summaries we compare are precisely the level at which large-scale fMRI organization is expressed, so this coarse-graining is well matched to the question we ask here, and in future studies. Second, we constrained connectivity using a single Euclidean distance kernel. This simplification omits white-matter connectivity, myelination, and cytoarchitectonic gradients, each of which also shapes cortical dynamics. How each of these constraints alters the correspondence between simulated and empirical dynamics is an open question. But, simple geometry is an established and reproducible predictor of cortical connectivity^1,2,18^, making it a principled and parsimonious starting point onto which richer constraints can later be layered. Third, although the DMS-D task engages working memory, temporal integration, perceptual decision making, and inhibitory control, it does not span the full range of cognitive demands; whether the same training trajectory holds across task families remains untested. Yet, it is notable that our *in silico bioRNNs* trained on one working-memory task predicted fMRI activity from a different working-memory task in humans, *in vivo*, suggesting that the correspondence may be anchored in shared cognitive demands rather than in superficial task features. It also suggests that future work that trains *bioRNNs* on multimodal tasks, *e.g.*, ones with simultaneous visual and auditory inputs, could further increase their biological relevance.

## CONCLUSION

Together, our findings demonstrate that physical geometry and cognitive inputs play distinct and complementary roles in the emergence of brain organization. While geometry constrains the space of possible dynamics, cognitive inputs determine which of those dynamics are expressed. Our findings further situate topology as the intermediate level of organization through which a geometrically constrained system reconciles the competing demands of wiring cost and computation, thereby enabling brain-like dynamics and cognitive performance to be realized simultaneously. By making the interactions among these three features observable and manipulable, *bioRNNs* turn long-standing questions about structure and function into direct investigations and offer a versatile *in silico* complement to human neuroimaging for investigating how the brain’s structure shapes its function. They additionally offer a potential avenue for performing individualized *in silico* counterfactual experiments that observational studies alone cannot achieve.

## METHODS

For the analyses described herein, we used coding assistance from a large language model (Claude Opus v4.6 and v4.8, Anthropic). We manually reviewed and modified all the code used in the analyses and assume full responsibility for the results.

### RNN overview

We compared three classes of recurrent neural network (RNN) that together form a graded hierarchy of biological constraint: *Vanilla RNNs*, *Masked RNNs*, and spatially embedded biophysical RNNs (*bioRNNs*). All three were instances of the same single-layer recurrent architecture and differed only in two configuration choices: whether their inputs and outputs were restricted to anatomically defined subsets of units (projection constraints), and whether their recurrent connectivity was shaped by a neurobiological spatial prior (brain geometry). *Vanilla RNNs* had neither constraint, *Masked RNNs* had projection constraints only, and *bioRNNs* had both projection constraints and a geometry-based prior on their recurrent weights. For each class, we trained 100 randomly initialized networks (runs). Because the classes were identical in every other respect, this design isolated the contribution of each constraint: the difference between *Vanilla* and *Masked* networks reflected the effect of projection constraints, whereas the difference between *Masked* and *bioRNNs* reflected the effect of the geometry prior alone.

### Network architecture

Each network consisted of 100 recurrently connected hidden units and was modeled as a discretized continuous-time (leaky) RNN. At each timestep, the hidden state, ℎ, evolved as

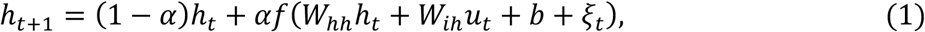

where *W_i_*_ℎ_ and *W*_ℎℎ_ are the input-to-hidden and hidden-to-hidden weight matrices, *u_t_* is the task input, *b* is a bias term, *f* is the hyperbolic-tangent nonlinearity, and *α* is the temporal integration factor. We set *α* = 0.1, resulting in a single-unit time constant 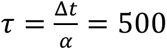 at an integration step of Δ*t* = 50 ms. The term *ξ_t_* is independent Gaussian recurrent noise added to the pre-activation hidden state during training only, with an effective standard deviation scaled as

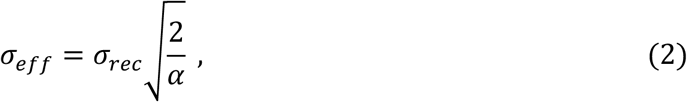

with *σ_rec_* = 0.05. We initialized the recurrent weights from a uniform distribution over the interval (−0.01, 0.01), and the remaining parameters used the default initialization. A linear readout mapped the hidden state to the task output at every timestep.

We placed the 100 hidden units in one-to-one correspondence with the 100 left-hemisphere parcels of the Schaefer 200-region, 7-network cortical atlas^33^. This mapping defined the spatial priors described below and was used to project network activity onto the cortical surface for comparison with empirical data.

### Projection constraints

In *Masked RNN* and *bioRNN* networks, task inputs entered the network only through a fixed subset of read-in nodes, and the decision was read out only from a separate subset of read-out nodes. Read-in nodes corresponded to the visual network (14 nodes), and read-out nodes corresponded to the default-mode network (27 nodes), following the Schaefer atlas network assignments. We enforced these constraints using binary masks: the input mask set the input weights of all non-visual nodes to zero, and the output mask set the readout weights of all non-default-mode nodes to zero. We reapplied the masks at every forward pass to prevent gradient updates from reintroducing masked connections. We left the recurrent weight matrix *W*_ℎℎ_ fully connected and did not subject it to these masks; projection constraints therefore restricted only which units received the stimulus and which drove the response, not the internal connectivity. *Vanilla RNNs* received inputs and produced outputs from all units, so they had no projection constraints.

### Spatial embedding and regularization

Left unconstrained, recurrent weights in RNNs can grow without bound and/or overfit. Thus, as is standard for RNN training, we regularized the hidden-to-hidden connectivity with a global L2 penalty added to the task loss. We used a variant of this penalty to constrain the network geometry^27^. The three RNN classes were subject to the same L2 penalty on the recurrent weights and only differed in whether that penalty incorporated cortical geometry. In all cases, a regularization term was added to the task loss and scaled by a fixed weight *λ* = 0.002.

For *Vanilla* and *Masked* networks, the recurrent weights penalty was a node-scaled L2 term,

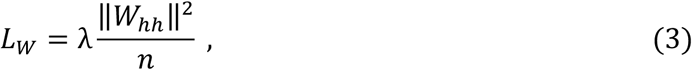

with *n* the number of nodes. This term applies a uniform weight penalty that contains no spatial information.

For *bioRNNs,* the penalty additionally aligned the recurrent connectivity with cortical geometry. We first computed the pairwise Euclidean distance matrix between the 100 left-hemisphere parcel centroids and normalized it by its mean off-diagonal value to obtain a distance kernel, *K*. We then defined a spatial proximity matrix, *S* = 1 − *K*. Thus, the *bioRNN* penalty combined a geometry term with the same scaled L2 decay according to

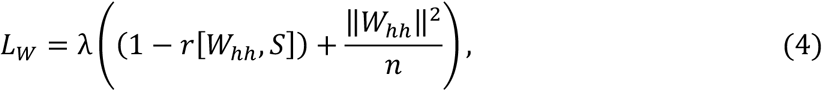

where *r*[*W*_ℎℎ_, *S*] is the Pearson correlation between the off-diagonal entries of the recurrent weight matrix and those of the similarity matrix. Minimizing this term drove the recurrent weights to co-vary with cortical proximity, so that nearby parcels in Euclidean space tended toward stronger coupling and more distant parcels toward weaker coupling. Because the scaled L2 term was identical across classes, the geometry term was the sole factor distinguishing *bioRNNs* from *Masked* networks.

### Task structure and training

We trained all networks on a delayed match-to-sample with distractors (DMS-D) task^31^ implemented in the NeuroGym toolbox^38^ and modeled after Yang et al.^32^. On each trial, the network first received a fixation input, followed by a sample stimulus, and then three test stimuli were presented in sequence, separated by delay periods (see example trial schematic in **Fig. 1d**). Stimuli were encoded across 32 input channels representing a one-dimensional circular feature, together with a fixation channel, giving a 33-dimensional observation. The sample and each test stimulus were presented for 500 ms, with 1,000 ms delays, at an integration step of Δ*_t_* = 50 ms. The network was required to produce a match response only when a test stimulus exactly matched the sample and to withhold responding otherwise. Successful performance, therefore, required maintaining the sample in working memory across the delays, comparing each test stimulus to it when presented, and inhibiting responses to non-matching (distractor) tests, thereby engaging working memory, temporal integration, perceptual decision-making, and inhibitory control.

### RNN training and learning-speed quantification

We trained the RNNs using gradient descent and the Adam optimizer^56^ with a learning rate of 0.001. We implemented all the RNNs in Python and trained them using PyTorch’s automatic differentiation. The task training objective (loss) was the cross-entropy between the per-timestep readout and the target action sequence. Thus, the loss that the RNNs were trained to minimize combined the task loss and recurrent weight regularization (spatial penalty) terms into a single overall loss,

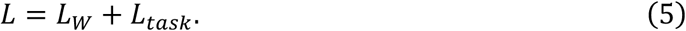

At each epoch, we produced a batch containing 32 sequences, which we split evenly into 16 training and 16 held-out validation sequences. Each sequence contained five consecutive task trials drawn from a per-run trial stream with a run-specific random seed. Initialization, trial stream, and recurrent-noise draws were determined by the run index, so that a given run index reproduced the same training conditions across classes and geometry permutations (see section on null testing), allowing us to isolate the observed effects to the spatial constraints. We trained the RNNs for 60,000 epochs and periodically checkpointed and evaluated their task accuracy on a fixed, independently seeded battery of 100 trials. A trial was scored as correct only when the network elicited the correct action within the permitted response window (*i.e.*, during the presentation of the matching test stimulus) and withheld responses otherwise. Training, validation, and test trials were drawn from disjoint random-seed ranges to prevent overlap.

We quantified RNN learning speed by fitting a logistic function to each run’s accuracy trajectory (**Fig. S1**), and defining the convergence epoch as the point at which the fitted curve reached 95% of that run’s asymptote. We then defined a per-RNN class training criterion as the 95th percentile of these per-run convergence epochs. We compared the three RNN classes using Wilcoxon signed-rank test with Holm-Bonferroni correction and report effect sizes using the matched-pairs rank-biserial correlation. We additionally established a global evaluation epoch to perform all subsequent analyses, defined as the epoch at which all RNNs across all classes had reached stable performance (asymptote). This resulted in epoch 40,000 being the global learning convergence point and was therefore the epoch at which we performed all analyses (except those that assessed the RNNs across epochs).

### Empirical fMRI data

We compared RNN dynamics against task functional MRI (fMRI) obtained from 100 typical young adults (50% male; 28.25 ± 3.67 years) from the Human Connectome Project^39^. Minimally pre-processed task fMRI time series were drawn from the working-memory N-back paradigm^43^. Time series were parcellated into the 100 left-hemisphere regions of the Schaefer-200, 7-network atlas to match the network nodes, and global signal regression was applied to attenuate widespread, non-neural fluctuations before the parcel-wise time series were extracted for comparison with the RNN dynamics. As a result, each subject’s task fMRI data was converted into a [time × parcels] matrix for subsequent analysis.

### Dimensionality reduction of network dynamics

To capture the dominant, behaviorally relevant modes of each network’s high-dimensional activity, we summarized hidden-state dynamics using principal component analysis (PCA)^40^. First, we tested each fully trained network on 100 task trials and concatenated the resulting hidden-state time series into a single [time × nodes] matrix. We ran PCA on this matrix using Scikit-learn’s^57^ PCA function with the exact solver, retaining the first five PCs, thereby extracting the low-dimensional, dominant modes of activity across nodes.

We considered two cognitive input regimes: a task regime (RNN_task_), in which the trained network was driven by the DMS-D task inputs, and a noise regime (RNN_noise_), in which it was driven by unstructured Gaussian noise (mean 0.5, standard deviation 0.3, with the fixation channel held fixed) in the absence of any task structure. The five-component PCA subspace obtained in each regime was used for the variance-explained analyses below.

### fMRI variance explained and the random-subspace baseline

We quantified how well each network’s PC subspace captured empirical brain dynamics as the fraction of a subject’s fMRI temporal variance that lies within that subspace. We centered each subject’s parcellated fMRI time series on its own temporal mean (per parcel), projected it onto the five-PC RNN subspace, and computed the fraction of temporal variance explained (VE). We did so for each combination of RNN class (n = 3), training run (n = 100), and subject (n = 100). Averaging this fraction across subjects gave the network’s VE for a single RNN run.

However, raw VE values cannot be interpreted directly because any five-dimensional subspace captures a non-trivial fraction of variance purely by virtue of its dimensionality^44^. To calibrate against this chance level, we established a random-subspace baseline. We drew 1,000 independent, uniformly random, five-dimensional orthonormal subspaces of a matched 100-dimensional space, each obtained as the orthonormal factor Q of the QR decomposition of a 100 × 5 matrix of i.i.d. standard-Gaussian entries. For every random subspace, we recomputed the identical subject-averaged VE metric, producing a null distribution of VE values expected in the absence of any genuine network–brain correspondence. We then expressed the network’s VE as a z-score relative to this null,

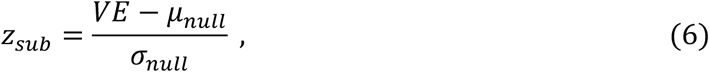

with *μ_null_* and *σ_null_* the mean and standard deviation of the 1,000 null VE values. Therefore, *z_sub_* reported, in standard-deviation units, how far a network’s PC subspace exceeded what an arbitrary subspace of matched dimensionality achieved: *z_sub_* ≈ 0 is chance-level correspondence, whereas a large positive *z_sub_* indicates that the specific directions spanned by the network’s dynamics, not simply their number, align with the structure of empirical fMRI. We applied the same approach when tracking *z_sub_* across the *bioRNN* training trajectory (hysteresis plot, **Fig. 3a**), and in the variance-decomposition analysis (**Fig. 3b**).

### Correspondence with the cortical hierarchy

We assessed whether the RNNs’ dynamics recapitulated the brain’s principal functional hierarchy, summarized by the sensorimotor–association (S–A) axis^47^. For each RNN class, we regressed the S–A axis values of the 100 left-hemisphere parcels onto the weights (spatial loadings) of the first five RNN PCs (averaged across RNN runs) and recorded the explained variance (adj. R^2^). For comparison, we performed the same regression using the PCs derived directly from the empirical N-back task fMRI data (averaged across subjects), providing an empirical benchmark for the extent to which low-dimensional dynamics align with the S–A axis.

### Geometry null testing

To test whether the *bioRNNs’* correspondence with empirical dynamics depended on the veridical anatomical arrangement of nodes rather than on the spatial autocorrelation that any distance-based kernel necessarily imposes, we constructed a geometry null. We generated spin permutations of the 100 left-hemisphere parcels by applying a uniformly random, three-dimensional rotation to their coordinates on the FreeSurfer spherical surface, and reassigning each rotated parcel to a distinct original parcel by optimal (Hungarian) bijective matching^13,46,58^. This procedure preserved the spatial autocorrelation of the embedding while breaking its correspondence to the exact cortical layout. We applied each resulting permutation as a distance kernel to fully retrain a *bioRNN* under the permuted geometry, while keeping the same task and projection constraints.

We trained 200 such permuted-geometry ensembles up to 40,000 epochs, the point at which networks had reliably reached the training criterion, and that served as the timepoint for all previous analyses. For every geometry (permutation), we used the same five run indices, and we chose those indices so that, in the veridical RNN, their VE values spanned the full range of the 100-run distribution. Because initialization, trial stream, and recurrent noise are all fixed by the run index (see *RNN training*), a given run index reproduces the identical training environment across every geometry, so that any difference between geometries is attributable to geometry alone. Therefore, we fully retrained 1,000 RNNs spanning 200 permuted geometries with five runs each.

In this design, VE could change for two reasons: the geometry itself and the run-to-run variability of training. Because our design was hierarchical, with multiple training runs nested within each permuted geometry, the per-run VE values were not independent, and pooling them would have conflated geometry effects with training variability, risking pseudo-replication^59^. Variance-components (linear mixed-effects) models are the standard approach for such nested data, partitioning variance into between-group (geometry) and within-group (training noise) components. Because the same five run indices recurred across all geometries, runs and geometries were crossed factors^60^; and because those indices were a small, deliberately chosen set rather than a random sample of runs, we modeled run as a fixed effect. We therefore fit a mixed-effects model using *statsmodels*^61^,

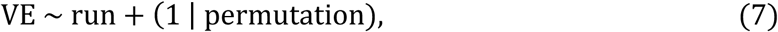

which estimated and removed the average effect of each run index (shared across geometries) before partitioning the remaining variance into a between-geometry component (*σ*_5*eom*_) and a within-geometry residual (*σ_train_*). Removing this deliberate run spread ensured that *σ*_5*eom*_ reflected only true geometry-to-geometry differences and *σ_train_* only residual training noise. We then asked whether the veridical geometry was an outlier relative to the distribution of permuted geometries, expressing its mean VE as a z-score against the between-geometry spread,

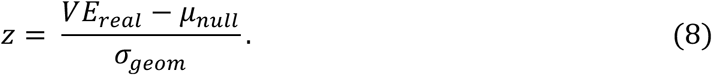

We converted this z-score into a one-sided p-value under the normal distribution; this parametric tail was justified because the distribution of permuted-geometry means did not deviate significantly from normality (Shapiro-Wilk *p* = 0.109). A cluster bootstrap (2,000 resamples over geometries and over the veridical RNN’s runs) then provided a 95% confidence interval on *z* (and, by association, on *p*). As an assumption-free check, the veridical geometry’s VE exceeded the Ves of the 200 permuted geometries (run-averaged).

### Training trajectory (hysteresis) analysis

To characterize how structure and function co-evolved during training, we tracked each *bioRNN* across 100 evenly spaced training checkpoints, in addition to the initial state (epoch zero). At each checkpoint, we computed two quantities: (i) structural similarity to the embedding, and (ii) fMRI variance explained. Structural similarity was calculated as the cosine similarity between the off-diagonal entries of *W*_ℎℎ_ and the corresponding entries of the spatial proximity matrix, *S*, computed over all 100 nodes. fMRI prediction was the fMRI variance-explained z-score (*z_sub_*) defined above, computed against a null of 1,000 random subspaces at each checkpoint. We also recorded task accuracy from the fixed test battery. We averaged these trajectories across the 100 runs and visualized fMRI prediction against structural similarity, coloring the trajectory by (log) task accuracy. Then, visually, we partitioned the trajectory into three phases: (i) the first phase spanned the RNNs’ initial rapid shift toward a high structural similarity value while its *z_sub_* remained below chance; (ii) the second phase was defined by the RNNs’ rapid shift toward a high *z_sub_* while its structural similarity to the embedding remained high; (iii) the final phase was defined as the one during which structural similarity began dropping and *z_sub_* remained high.

### Variance decomposition of task-evoked and intrinsic dynamics

To isolate the contribution of structured task input to the *bioRNN* predictive dynamics, we compared fully trained *bioRNNs* driven by task inputs against the same networks driven by unstructured noise (the task and noise regimes above). We applied a commonality analysis, a variance decomposition method that separates the variance explained uniquely by each predictor set from the variance they explain jointly^62,63^. For each regime, we obtained a five-dimensional PC loading matrix (as we did in the initial fMRI prediction analysis) and formed a pooled basis by performing a singular value decomposition of the concatenated task and noise loadings (resulting in a basis of up to 10 dimensions). The task-only and noise-only subspaces each define the directions in fMRI space that one input regime can explain, whereas the pooled basis represents everything either regime can explain; therefore, contrasting the pooled fit against each regime-specific fit isolates the variance that is unique to task input, unique to noise input, or shared between them. For each subject and each RNN run, we computed the fraction of variance in that subject’s fMRI time series captured by the task-only subspace (*VE_task_*), the noise-only subspace (*VE_noise_*), and the pooled subspace (*VE_pooled_*). We then partitioned these into a unique-task component (*VE_task_*__*only*_), a unique-noise component (*VE_noise_*__*only*_), and a shared component (*VE_s_*_ℎ*ared*_):

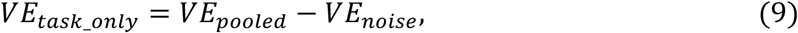

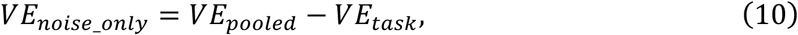

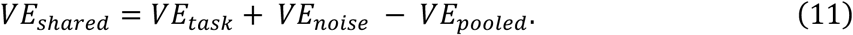

Again, significance was assessed against a null of 1,000 random orthonormal subspaces.

### Emergent network topology

We characterized the graph-theoretic organization of the intermediate nodes, *i.e.*, those designated as neither read-in nor read-out, and therefore outside the direct influence of the *projection constraints*. From the recurrent weight submatrix among these units, we built a graph by taking the absolute value of each connection, zeroing the diagonal, and retaining only the strongest 20% of edges. Because the recurrent weights are directed, the resulting graph is also weighted and directed; we therefore performed the analyses with directed, weighted graph metrics computed using the Python implementation of the Brain Connectivity Toolbox (*bctpy*)^64^.

We quantified five aspects of network organization. To index the emergence of hubs, we computed each node’s total (in-plus-out) degree (*bctpy*: *degrees_dir*) and took the mean degree of the top 20% highest-degree nodes (**Fig. 4a**). To test whether those hubs formed an interconnected core, we computed the weighted directed rich-club coefficient (*bctpy*: *rich_club_wd*) and normalized it against an ensemble of 1,000 degree-preserving randomized graphs (*bctpy*: *randmio_dir*), following the normalized rich-club approach of van den Heuvel & Sporns^17^ (**Fig. 4b**). To measure local segregation, we computed the mean weighted directed clustering coefficient (*bctpy*: *clustering_coef_wd*)^65^ (**Fig. 4c**). To measure cross-module integration, we first partitioned the graph into communities by directed modularity maximization (*bctpy*: *modularity_dir*)^66^, then computed the mean participation coefficient across nodes with respect to that partition (*bctpy*: *participation_coef*)^67^ (**Fig. 4d**). The fifth metric captured the physical cost of connectivity; we computed the mean connection length of the retained (top 20%) edges, defined as the mean Euclidean distance between the parcel centroids that each edge connects, using the same inter-regional distance matrix that defined the spatial embedding (**Fig. 4e**).

We evaluated all five metrics at each training checkpoint spanning the third phase of the hysteresis trajectory (see *Training trajectory analysis*). To summarize the timing of topological change, we rescaled each metric by min-max normalizing its values across training to the unit interval, so that metrics with different ranges could be compared on a common scale, and then calculated their temporal derivative with respect to training epoch (**Fig. 4f–j**).

### Statistical analysis and reproducibility

Plots are generally generated using mean values for group summaries and 95% confidence intervals, computed as 1.96 times the standard error of the mean, as shaded bands. For values reported in the text, unless otherwise stated, we report the mean ± SD.

## DATA AND CODE AVAILABILITY

All RNNs were implemented in Python (v3.12.11) using PyTorch (v2.7.1)^68^. The RNNs were implemented as a custom module based on the PyTorch RNN class and optimized using PyTorch’s automatic differentiation engine (see *RNN training*). The DMS-D task was simulated using NeuroGym (v2.1.0)^38^. Training was run on a Mac Studio with the M3 Ultra chip. The code used in this paper for RNN training and all reported analyses is available in the *neuro_rnn* GitHub repository (https://github.com/LindenParkesLab/neuro_rnn). The human fMRI data is part of the publicly available Human Connectome Project.

## ACKNOWLEDGEMENTS

LP was supported by the National Institute Of Mental Health of the National Institutes of Health under Award Number R00MH127296. Research reported in this publication was also supported by Rutgers University through the Rutgers Health BMIHAI PAIR Postdoctoral Fellowship. WUB received partial support from a seed project funded by the Office of the Vice Provost for Research at Rutgers University. The content is solely the responsibility of the authors and does not necessarily represent the official views of the institution. Data were provided in part by the Human Connectome Project, WU-Minn Consortium (Principal Investigators: David Van Essen and Kamil Ugurbil; 1U54MH091657) funded by the 16 NIH Institutes and Centers that support the NIH Blueprint for Neuroscience Research; and by the McDonnell Center for Systems Neuroscience at Washington University.

