## Supplementary Materials for "Geometric constraints and cognitive inputs jointly shape emergent brain dynamics and topology"

Ahmad Beyh [1], Jason Z. Kim [2], Waheed U. Bajwa [3], Linden Parkes [1,4]

[1] *Department of Psychiatry, Brain Health Institute, Rutgers University, Piscataway, NJ, USA*
[2] *Department of Physics, Cornell University, Ithaca, NY, USA*
[3] *Departments of Electrical & Computer Engineering and Statistics, Rutgers University, Piscataway, NJ, USA*

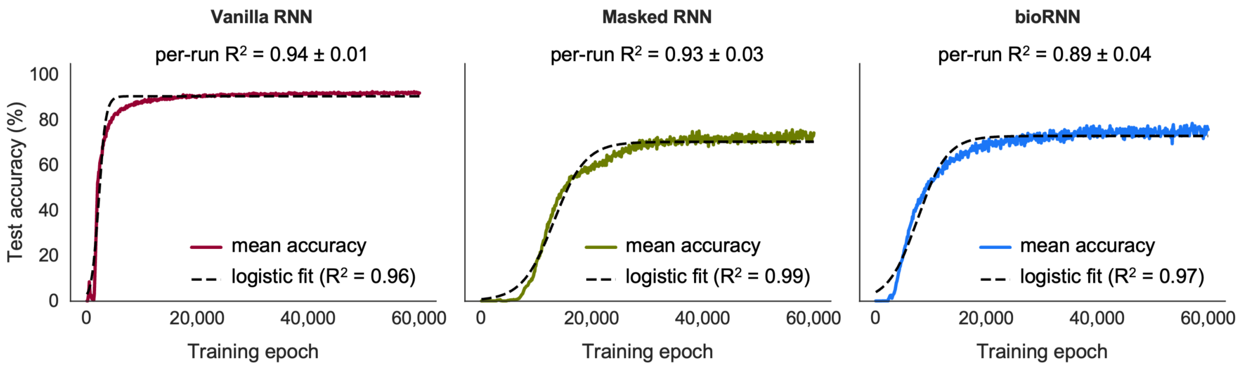


**Figure S1. Task performance sigmoid fits.**
We modeled the task performance curve (solid line) of each model as a logistic function (dashed line) and used it to quantify training convergence. The sigmoid was a suitable model, as shown by the $R^2$ values: 0.96 (*Vanilla RNN*), 0.99 (*Masked RNN*), and 0.97 (*bioRNN*).

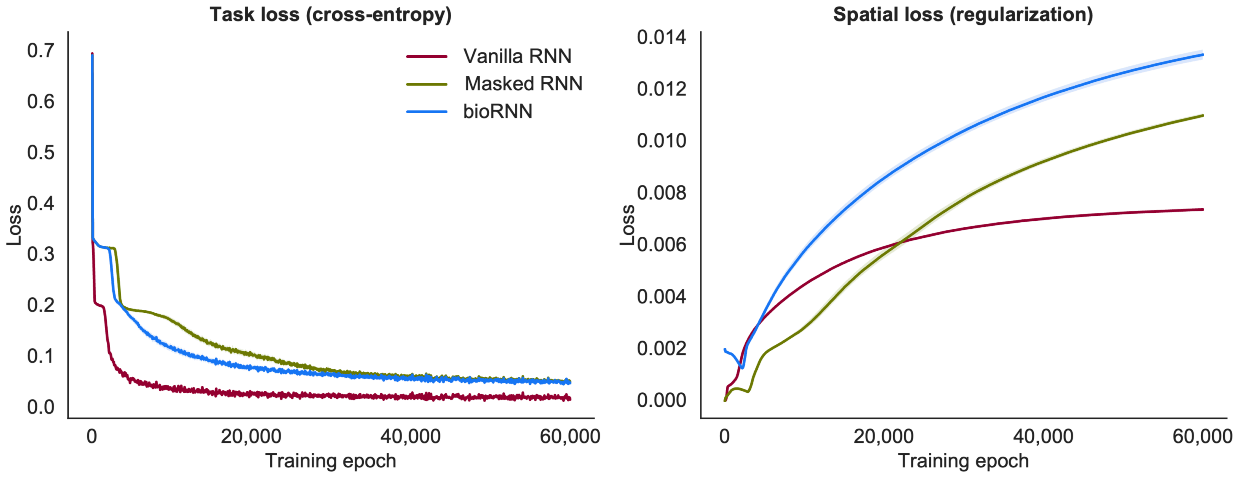


**Figure S2. Spatial and task loss terms across model training.**
The spatial loss (L2 only for *Vanilla RNNs* and *Masked RNNs*, and L2+embedding for *bioRNNs*) was the highest, *i.e.*, imposed the strongest penalty for *bioRNNs*. However, task loss (cross entropy) was lower for *bioRNNs* compared to *Masked RNNs*. This indicates that, despite imposing an additional constraint during training, the spatial embedding allowed *bioRNNs* to learn the task better than *Masked RNNs*.

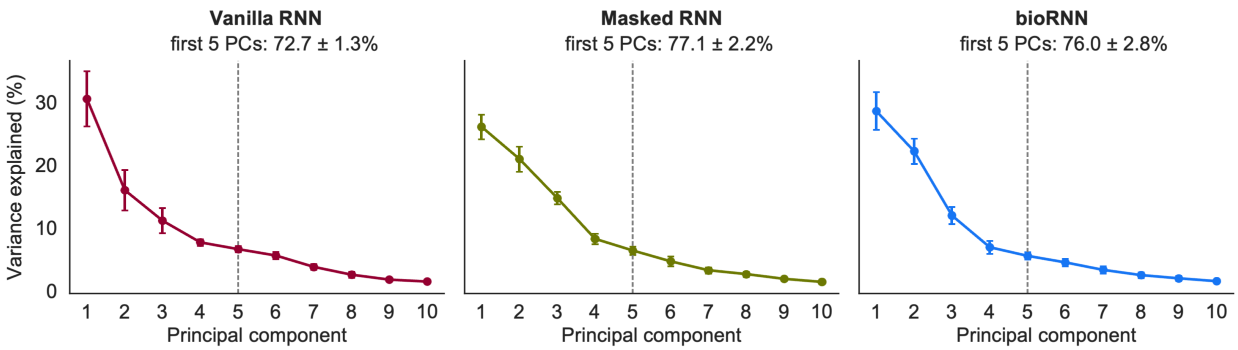


**Figure S3. PCA explained variance in the trained RNNs.**
Each plot shows the mean explained variance by the first 10 PCs for each RNN class. The error bars represent the standard deviation across 100 runs per class. We retained the first five PCs (vertical dashed lines) for the main analysis, as this included the inflection point in explained variance for all three classes.

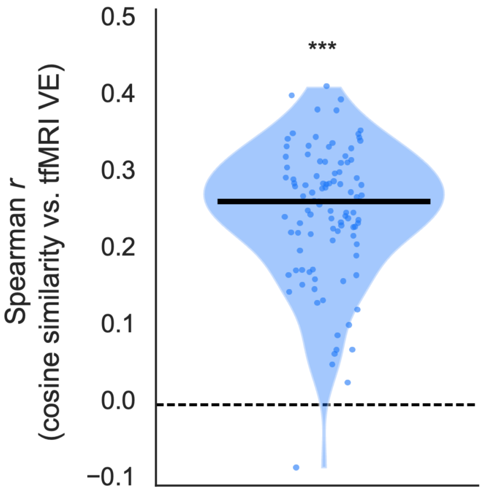


**Figure S4. Subject-specific effects.**
For each subject, we assessed how well the similarity of a *bioRNN*'s hidden weights to that subject's Euclidean geometry correlated with the ability of the *bioRNN* to predict that subject's fMRI dynamics. We observed a consistent positive correlation: $r_S = 0.25 \pm 0.09$, $p < 0.001$. Each dot represents the correlation between 100 *bioRNN* runs and one subject.

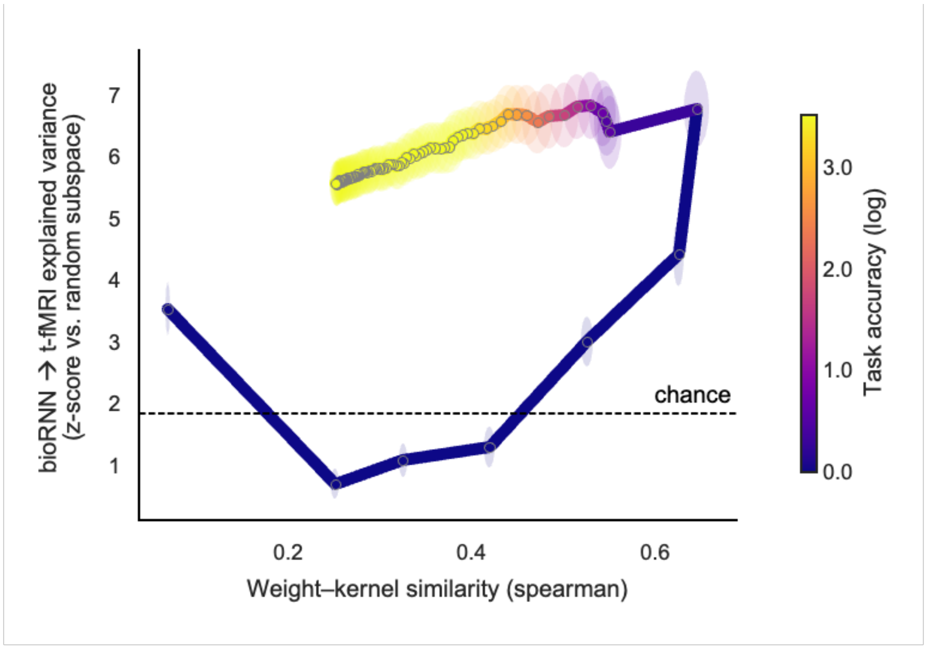


**Figure S5. Hysteresis loop trajectory under a Spearman similarity metric.**
We reproduced the hysteresis loop effect that we observed in the main analysis (**Fig. 3**) using a Spearman correlation metric to assess the similarity between the *bioRNN* hidden weights matrix and the Euclidean embedding matrix. We observed that the same effect persists, indicating that the trajectory is not simply a product of the similarity metric.

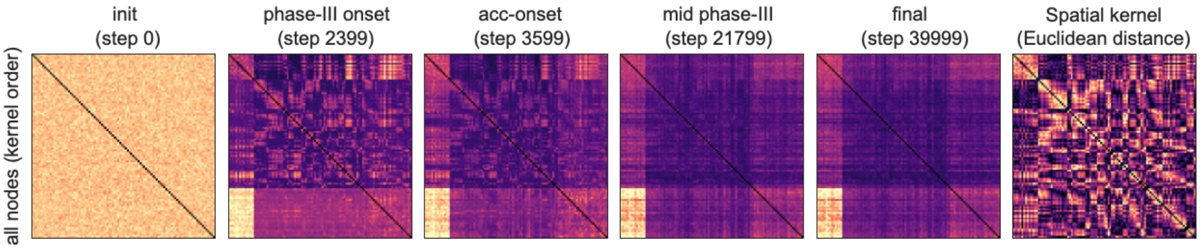


**Figure S6. Snapshots of the bioRNN hidden weights across training.**
Because of the projection constraints we impose on the networks during training, we expected that the *bioRNNs* would develop a 'tripartite' overall topology. These snapshots of the models' hidden weights at various points along the training timeline showcase this and are the motivation for using the 'bystander' nodes only (*i.e.*, non-input and non-output nodes) for the topology analysis. For comparison, the last panel shows the Euclidean distance embedding kernel used to train the models.

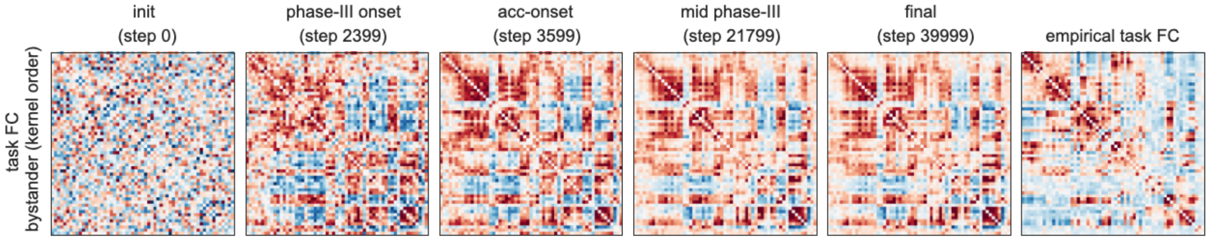


**Figure S7. Functional connectivity in the bioRNN model across training.**
The 'bystander' subnetwork in the *bioRNNs* developed a functional connectivity pattern suggesting the presence of modules and exhibited a qualitatively similar pattern to that of empirical (*in vivo*) functional connectivity observed in 100 HCP subjects (rightmost panel). Importantly, these are the nodes that were not subject to *projection constraints* and were only shaped by task learning and the spatial embedding.

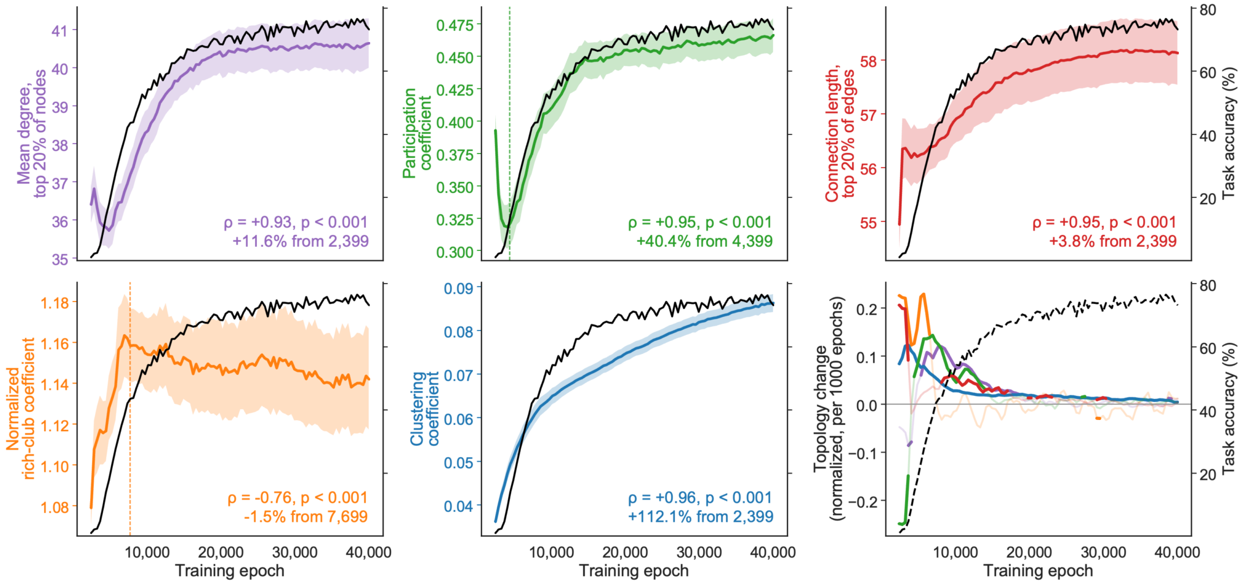


**Figure S8. Topology in *bioRNNs*.**

*bioRNNs* developed features of brain topology. The networks developed hubs that were selectively inter-connected (high degree and rich-club >1), as well as inter-connected modules (clustering and participation). In addition, the networks selectively increased the weights of long-range connections. These features developed rapidly just before RNNs solved the DMS-D task and then became increasingly stable as performance plateaued. Where present, the dashed vertical line indicates the start of the window from which we derive the displayed statistics. This was done to avoid the undesired effects of the initial large fluctuations.

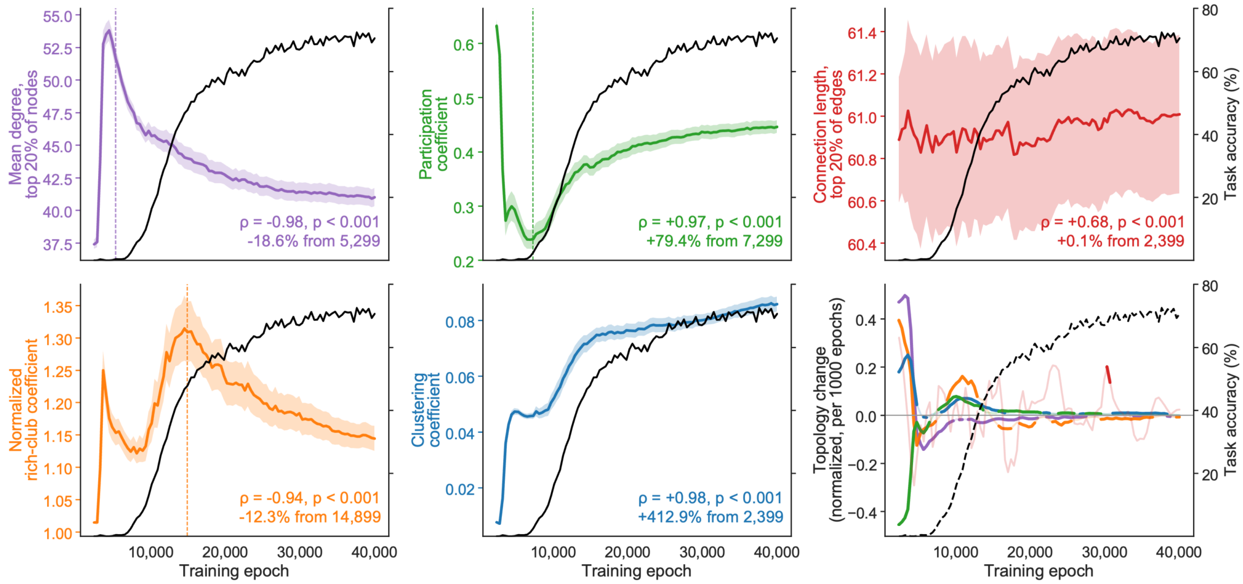


**Figure S9. Topology in *Masked RNNs*.**

The *Masked RNNs* developed some features of brain topology, such as clustering and participation, but other features, such as hubs and rich-club, emerged early but were not maintained, as evidenced by their sharp decline as the RNNs improved task performance. In addition, the models did not develop any specific pathways that mapped selectively onto long-range connections, as *bioRNNs* did, but this was not expected, given that these models were not exposed to the Euclidean embedding kernel. Where present, the dashed vertical line indicates the start of the window from which we derive the displayed statistics. This was done to avoid the undesired effects of the initial large fluctuations.

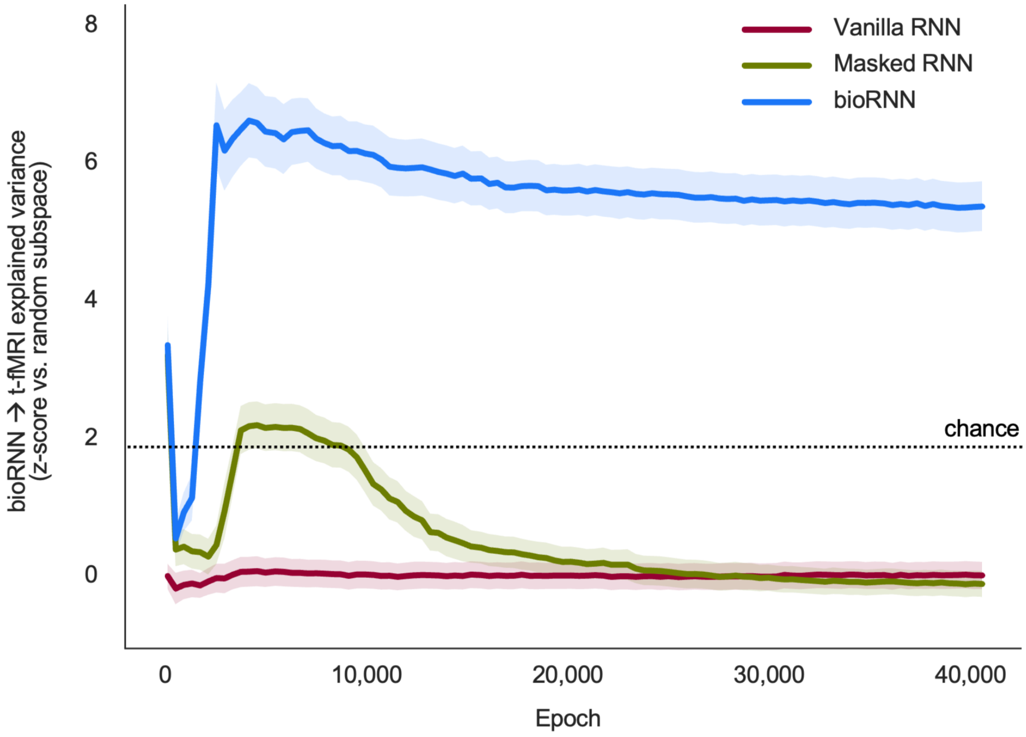


**Figure S10. Timeline of RNN prediction of fMRI dynamics.**
The three RNN classes we trained had markedly different abilities to predict fMRI activity throughout training. *Vanilla RNNs* remained very poor predictors throughout the training process. *Masked RNNs* seemed to develop some ability to predict fMRI activity, but this early 'bump' faded as the networks improved their task performance. This also coincided with a drop in certain topological features (**Fig. S9**). Meanwhile, *bioRNNs* were the only models to remain highly predictive throughout most of their training trajectories, and this was accompanied by the emergence of several brain-like topological features (**Fig. 4** and **Fig. S8**).
